# Network-Based Kidney Allocation Simulation: Evaluating Organ Matching Strategies in Variable Hospital Networks

**DOI:** 10.1101/2025.04.22.650043

**Authors:** Aniruth Ananthananarayanan, Benjamin Hu, Alex Sha

**Affiliations:** University of North Texas Denton, TX

## Abstract

Kidney allocation is a complex and high-stakes process shaped by various factors. Although existing simulations have improved our understanding of transplant policies, they often stem from the perspectives of medical urgency and biological compatibility. Hence, they often fail to account for the geographic and structural variability of hospital networks that heavily influence organ distribution. In this paper, we present a modular network-based kidney allocation simulation designed to function as a standardized benchmark to evaluate organ matching strategies in more realistic hospital networks. Our simulation models hospitals as nodes in a dynamic network that incorporates detailed donor-recipient compatibility criteria and provides an intuitive interface for testing custom matching policies. Then, through two case studies, we demonstrate how varying network structures and matching strategies affect patient outcomes, quantified through transplant success rates and the percentage of positive patient outcomes. Through these case studies, we derive results that closely resemble real-world observations and identify clear biological explanations for the observed trends, aligning with established empirical knowledge. Our results highlight the importance of geography-/network-aware benchmarks and allocation strategies and provide a standardized platform for future research on optimizing kidney distribution in healthcare networks.

## 1 Introduction

The allocation of donated kidneys represents a significant challenge in organ transplant policy [1]. The scarcity of available kidneys, combined with the ever-growing demand for transplants, has driven the development of various frameworks that aim to maximize patient outcomes. These frameworks are based on theoretical models and are designed to optimize the success rate based on factors such as patient survival, graft success, and overall quality of life. However, testing these models in real-world clinical settings comes with profound ethical and practical risks: mistakes in allocation could directly compromise patient lives and waste valuable organs.

Simulation methods allow researchers to assess the impact of new allocation strategies without endangering patients. For example, frameworks such as simKAP and SRTR’s Kidney Allocation Policy Simulation Allocation Model (KPSAM) have advanced our understanding of clinical decision-making and the dynamics of patient waiting lists [2, 3]. Despite these advances, existing simulation methods often neglect the role of geographic network structures that are fundamental to real-world allocation; the geographic spread of hospitals can significantly influence both the accessibility and efficiency of kidney distribution [4, 5].

To bridge this gap, we have developed a comprehensive simulation that directly incorporates hospital networks and inter-hospital organ sharing dynamics. Our aim is to establish a standardized benchmark that more accurately reflects the complexities encountered in real-world scenarios [6].

## 2 Benchmark Construction

### 2.1 Network Simulation Design

Our framework represents the kidney allocation system as a network of connected hospitals and transplant centers; nodes represent individual hospitals/transplant centers and edges represent organ transfer routes with associated travel times. This approach enables the evaluation of how network topology influences allocation efficiency with patient outcomes.

### 2.2 System Architecture

The kidney transplant simulation system is composed of several interconnected classes that model various aspects of the organ allocation process. These include the `Donor, Organ`, and `Patient` classes, which represent the basic entities within the system, and the `Hospital` and `TransplantCenter` classes that simulate the operation of hospitals and the broader hospital network. A high-level overview of the system is as follows:

#### 1. Domain Classes

- **Donor Class:** This class models organ donors, including donor attributes such as blood type, age, HLA (Human Leukocyte Antigen) profile, and donation type (living or deceased). These factors are crucial in determining the compatibility between donors and recipients.
- **Organ Class:** This class represents a donated kidney and includes attributes such as preservation method, transportation logistics, and quality metrics (e.g., expected lifespan and condition).
- **Patient Class:** This class models transplant candidates, with key attributes including blood type, HLA, urgency level, and PRA (Panel Reactive Antibody), which influence their eligibility for a transplant.

#### 2. Hospital and Network Structure

- **Hospital Class:** The class maintains a list of patients awaiting transplants and an inventory of available organs. The hospitals perform the matching process and record the results of each transplant.
- **TransplantCenter Class:** This class represents a network of interconnected hospitals. The class coordinates organ sharing and ensures efficient organ distribution across the network, accounting for factors such as travel time between hospitals.

#### 3. Simulation Framework

- The `Simulation` class is responsible for managing the overall simulation process. The class manages the flow of time, updates patient waiting times, introduces new patients and organs into the system, and orchestrates the matching and transplant processes. Simulation parameters, such as organ and patient arrival probabilities, can be configured based on the user’s requirements.

### 2.3 Simulation Process

At each simulation step, the system updates the status of all patients on the waiting list and introduces new patients and organs to the network based on pre-defined probabilities. The following sequence occurs during each step:

- **Patient and Organ Arrival:** New patients and donated organs are introduced to the hospitals according to arrival probabilities, ensuring that the system models a dynamic, real-world environment.
- **Matching Process:** The system performs a matching procedure across the hospital network, selecting the most compatible organ for each patient. Each match is evaluated based on the factors listed in the previous section.
- **Transplant Outcome Simulation:** Once a match is made, a transplant is simulated, and the outcome (successful transplant or rejection) is determined based on the matching score and simulated success probability.
- **Metrics Calculation:** After each transplant, the system calculates detailed metrics, including transplant success rates, average waiting times, matching scores, and organ quality breakdowns.

### 2.4 Other Features

The simulation system also includes several other features to enhance its functionality:

- **Parameter Sweeping:** The system supports the execution of multiple simulations with varying parameters (e.g., organ/patient arrival probabilities, matching criteria) to optimize kidney allocation strategies.
- **Customizable Matching Strategies:** Users have the flexibility to implement custom matching algorithms as functions in Python, allowing for the exploration of different approaches to organ allocation.
- **Detailed Logging:** Comprehensive logs are maintained throughout the simulation, recording every matching decision, transplant outcome, and other key events. This data is invaluable for post-simulation analysis and optimization.
- **Multi-threading:** Parameter sweeps are routed through a thread pool executor for high-throughput concurrent simulations.

By incorporating these components and features, the simulation provides a flexible and realistic platform for analyzing kidney allocation policies and testing new strategies under various conditions. The system’s modular design also allows for easy extension, enabling users to customize the simulation to fit their specific research needs.

## 3 Experiments

### 3.1 Matching Policy Implementations

To further explore the motivation behind a standardized benchmark, we design two case studies that compare 4 different simplistic matching policies.

- **Default Strategy:** This policy uses the patient’s built-in compatibility calculation without any modifications. It serves as the baseline for comparison.
- **Wait Time Emphasis:** This policy prioritizes patients who have been on the waiting list for a longer period. A bonus is added to the compatibility score based on the patient’s wait time, ensuring that long-waiting patients have an improved chance of receiving a transplant.
- **Urgency Emphasis:** This strategy prioritizes patients with higher medical urgency by adding a significant bonus to the compatibility score proportional to urgency. The goal is to ensure that critically ill patients receive transplants sooner.
- **HLA Emphasis:** This approach places greater importance on human leukocyte antigen compatibility (HLA) between the donor and recipient. The matching score assigns a higher weight to HLA match scores while also accounting for donor-patient size compatibility and organ quality. This method aims to improve long-term transplant success rates.

Each of these strategies represents a different approach to kidney allocation, allowing for a comparative analysis of their effectiveness in terms of transplant success, waiting times, and overall patient outcomes.

### 3.2 Case Study 1: Comparative Algorithm Analysis

This case study evaluates the four matching policies by running simulations under identical conditions. The goal is to assess how different matching strategies impact key performance indicators–such as the percentage of transplants that were successful–and understand how changes in environmental conditions–such as network size, organ arrival probability, etc.–affect the policy’s effectiveness.

More specifically, we vary the organ arrival probability, patient arrival probability, and number of nodes in the network. We measure each policy’s effectiveness by tracking the number of successful transplants and patient outcomes over multiple simulation runs.

### 3.3 Case Study 2: Effects of Network Composition

This case study explores how variations in hospital network structure impact matching outcomes. Specifically, we examine how the ratio of transplant centers to hospitals, which we term the network composition, influences kidney allocation and whether certain policies perform better in specific network compositions.

To ensure a fair comparison, we maintain a constant total organ arrival rate across all network configurations. This means that the product of the organ arrival probability per transplant center and the number of centers remains fixed. By doing so, we isolate the effect of network composition on transplant success rates and patient wait times, allowing us to assess how different policies perform under varying levels of decentralization.

We measure each policy’s effectiveness by tracking the number of successful transplants and the average patient wait time over multiple simulation runs.

## 4 Results and Discussion

Kidney transplantation success is determined by a variety of factors such as donor and recipient HLA compatibility, ischemic time of the organ, and patient health prior to the transplant. To maximize patient outcomes, kidney allocation networks must carefully balance these variables. Our novel simulation model was designed with these traditional constraints in mind while also taking into account the role of geographic network structures in kidney allocation. Across both case studies, our simulation model reveals that strategies that emphasize HLA compatibility tend to perform significantly worse than the default strategy and those that prioritize urgency and wait-time. Additionally, our model shows how changes in organ arrival rates and network structure impact transplant success rates in a way that is consistent with existing practices. These results validate the relevance of our model and suggest its usefulness as a tool for evaluating the effectiveness of future kidney transplantation policies.

One key insight we have observed from our simulation involves the relationship between organ arrival probability and transplant success rates. As shown in Figure 1, our model demonstrates that as organ availability increases, the success rate of transplants decreases across all matching strategies. When organ arrival probabilities are low, hospitals and transplant centers tend to prioritize higher quality matches, focusing on maximizing donor-recipient HLA compatibility and minimizing ischemic time. However, as organs become more readily available, allocation strategies shift from finding the “perfect” recipient for each organ to efficiently making a greater number of “good” matches. This approach allows a larger number of patients to receive transplants; however, the overall transplant success rate slightly declines due to an increased likelihood of suboptimal biological matches. Our simulation captures this tradeoff, emphasizing its usefulness as a tool for researchers to optimize allocation strategies under real-world constraints.

**Figure 1.**
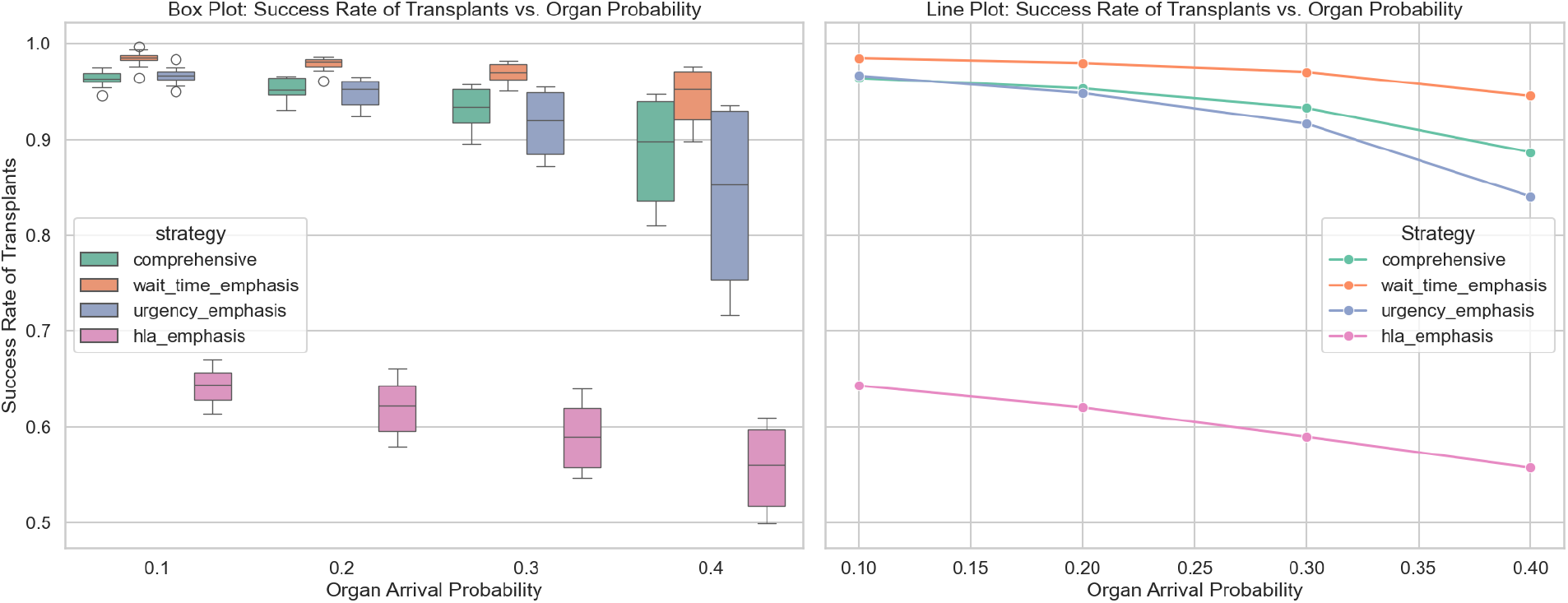
A comparison of the four core organ management metrics we defined in earlier sections. Here, we define the success rate as the number of successful transplants divided by the total number of transplants that occurred. Measurements were taken over a sample size of 100 simulations with each configuration.

Our simulation results also align with findings from recent studies regarding the relationship between prioritizing donor-recipient HLA compatibility and minimizing patient wait times. Figures 1 and 2 both clearly illustrate that the HLA-emphasis model consistently performs significantly worse than the other three models we tested in terms of transplant success rates and average wait time. This is a result of the model’s strict prioritization of donor-recipient HLA compatibility, which increases patient wait times by delaying matching until a highly compatible donor is located [7, 8].

**Figure 2.**
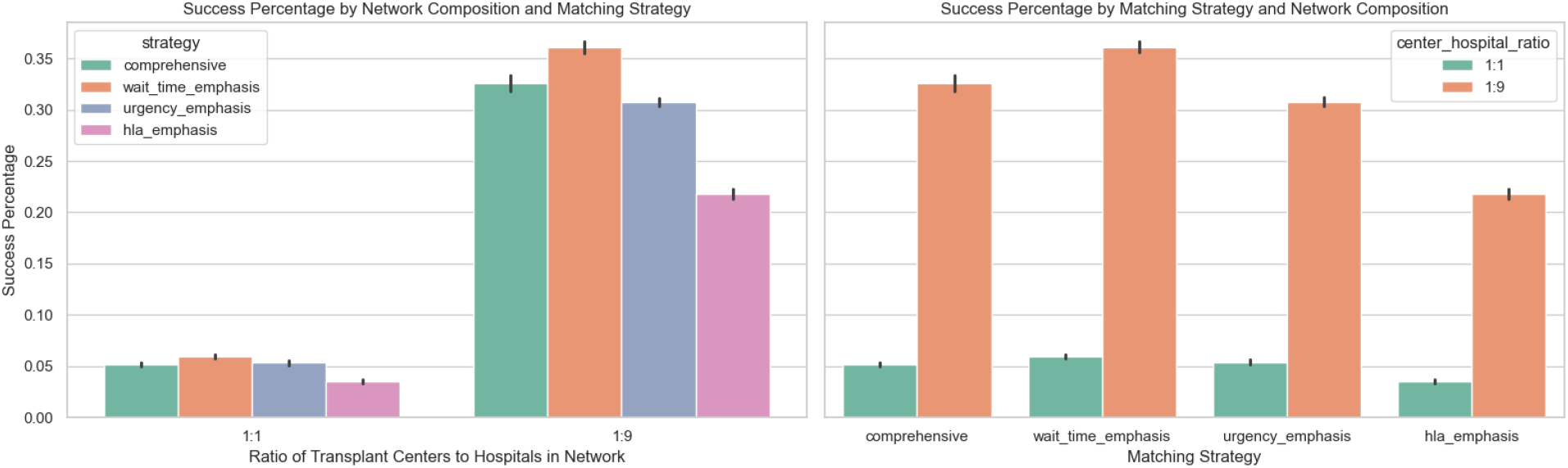
Results compare four strategies across three center-to-hospital ratios. HLA emphasis underperforms, while urgency and wait time strategies improve outcomes, especially in centralized networks. Measurements were taken over a sample size of 100 simulations with each configuration, and error bars show one standard deviation.

Our simulation also revealed another clear trend: networks with fewer transplant centers correlate with a higher transplant success percentage and lower wait times. Although it may seem that more transplant centers would suggest broader access, the concentration of resources and expertise in fewer, high-volume centers appears to streamline the transplant process and improve patient outcomes. High-volume institutions like the Mayo Clinic and the Cleveland Clinic report significantly better post-transplant survival rates, which have been largely attributed to standardized surgical protocols, experienced medical teams, and collaboration across multiple departments [9, 10]. Additionally, fewer transplant centers allow for more efficient organ distribution. In a network with many transplant centers, organs must be matched and transported across a more complex and fragmented network, often requiring longer travel distances and coordination between multiple facilities, increasing the time from donation to transplantation. Conversely, in a network with high-volume transplant centers, patients experience shorter wait times because they have access to a larger donor pool, organs are transported more efficiently, and there are fewer logistical redundancies within the system [11].

After analyzing our simulation results, it is clear that traditional organ transplant strategies that prioritize HLA compatibility are outperformed by those that emphasize patient urgency and waiting time. As such, it is necessary for real-world kidney allocation systems to continue to shift away from rigid matching models that focus on finding the perfect biological recipient for each organ to more flexible approaches that emphasize reducing patient waiting time. Given the adverse health effects of prolonged dialysis, transplants are much more likely to be successful when patients receive a moderately matched kidney sooner than waiting for a perfect match [12]. Due to modern advances in immunosuppressive therapy, patients today can tolerate less-than-perfect HLA matches without significantly increasing the risk of rejection [13].

Additionally, policymakers should consider optimizing the distribution of transplant centers to streamline the organ matching and distribution system. Our simulation has demonstrated that fewer, high-volume transplant centers lead to better patient outcomes, primarily by reducing patient wait times. Not only does centralizing resources into a smaller number of facilities increase organ allocation efficiency, reducing ischemic time, but it also ensures that surgical teams have more experience with complex transplant procedures. Rather than spreading already limited resources thinly across many facilities, our model and real-world studies confirm that concentrating organs in specialized facilities maximizes both organ distribution efficiency rates and expands patient access to a broader donor pool [14].

## Code Availability

The Python code for the benchmark is published on Github at https://github.com/AniruthAnanth/kidneybench.

